# SNP heritability: What are we estimating?

**DOI:** 10.1101/2020.09.15.276121

**Authors:** Konrad Rawlik, Oriol Canela-Xandri, John Woolliams, Albert Tenesa

**Affiliations:** The Roslin Institute, Royal (Dick) School of Veterinary Studies, The University of Edinburgh, Easter Bush Campus, Midlothian, EH25 9RG. Scotland. UK; MRC HGU at the MRC IGMM, University of Edinburgh, Western General Hospital, Crewe Road South, Edinburgh. EH4 2XU. UK

## Abstract

The SNP heritability 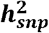 has become a central concept in the study of complex traits. Estimation of 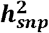 based on genomic variance components in a linear mixed model using restricted maximum likelihood has been widely adopted as the method of choice were individual level data are available. Empirical results have suggested that this approach is not robust if the population of interest departs from the assumed statistical model. Prolonged debate of the appropriate model choice has yielded a number of approaches to account for frequency- and linkage disequilibrium dependent genetic architectures. Here we analytically resolve the question of how these estimates relate to 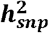 of the population from which samples are drawn. In particular, we show that the correct model for the purpose of inference about 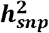 does not require knowledge of the true genetic architecture of a trait. More generally, our results provide a complete perspective of these class of estimators of 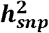, highlighting practical shortcomings of current practise. We illustrate our theoretical results using simulations and data from UK Biobank.

---

The SNP heritability is defined as the fraction of the phenotypic variance explained by additive effects of a given set of genetic variants^1,2^. It forms a bound on the ability to predict a phenotype using linear models of the chosen variants and has been important in the debate about the so-called missing heritability^3–5^. It has also been used to draw conclusions about the genetic architecture of phenotypes by contrasting the heritabilities of different categories of genetic variants^6^. Variance components, based on genomic relationship matrices, fitted using restricted maximum likelihood estimation, the so called G-REML method^3^, have been proposed, implemented in various tools^7–10^, and widely adopted as an approach to estimate 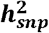. However, typically the quantity of interest is 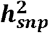 for the wider population from which samples were obtained. This population is not sampled from the statistical model underlying variance component estimation. It is therefore unclear to what extent the parameters estimated using the assumed model can be related to the quantity of interest, namely parameters of the wider population from which samples were obtained. This is primarily the case because the estimates are not directly available in analytical form, but are obtained using a numerical optimization procedure. Moreover it is known that G-REML cannot estimate 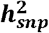 in general, as the estimates are, unlike 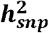, not invariant under general linear transformations of the genotypes^1^.

Based on empirical observations, it has been suggested that G-REML estimates are biased under departure of the population from the model^4,8,11,12^. This has led to a number of variations on the G-REML approach based on various assumptions about the genetic architecture of the phenotype^8,11–13^. These different models yield different estimates^14,15^. However, the correct choice of model remains unclear, as does the question what aspects of the population need to be incorporated into the model. We show that these questions can be resolved analytically, by directly relating the estimates of model parameters to parameters of the population. Our results highlight the central role of the linkage disequilibrium (LD) structure of the population, in particular between variants with non-zero additive effect. We show that incorporating this structure, which can be estimated from data, into the genomic relationship matrix (GRM) leads to an statistically consistent estimator of 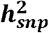. Conversely, the choice of alternative GRMs leads to a bias. This bias depends on the departure of the GRM’s assumptions from the LD structure of the population. It furthermore changes depending on the relative number of individuals and genetic variants in the analysis. The G-REML model with the standard GRM corresponds to the assumption of perfect linkage equilibrium amongst all modeled genetic variants. In practice populations are expected to depart from this assumption. On the one hand, natural processes like assortative mating or selection act directly on the LD structure in the population^16^. On the other hand, sampling strategies like, for example, case-control sampling will induce LD between causal variants in the sample^17^. Finally, we show that these effects are relevant even if causal variants are in linkage equilibrium. Typically, the set of modeled variants will not include all causal variants. We show, that this is sufficient to necessitate accounting for the LD structure of the modeled variants in order to avoid complex biases.

## Results

### Relating G-REML estimates to population parameters

The G-REML estimate of 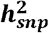, is given by 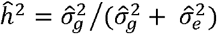 where 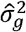, 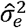 are the restricted maximum likelihood estimates of genetic and environmental variance components (Methods). Here we will concentrate on the more interesting estimate of 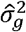. Furthermore, we will restrict our discussion to the form of the results in a setting which allows easier interpretation. Amongst others, we will assume that no fixed effects beyond the mean are fitted in the model and sufficiently large sample sizes. These and other implicit assumptions made here can however be relaxed, yielding a more generally applicable form of the main result (Methods). The genetic variance component is modeled using a GRM computed from the genotypes at the modeled genetic variants. Commonly employed GRMs are of the form **G = ZΣZ**^*T*^, where **Z** is the *N* × *M* matrix of standardized, i.e., centered and unit variance scaled, genotypes for *M* genetic variants of *N* individuals and **Σ** is a matrix which differs between GRMs (Methods). We show that 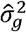 satisfies

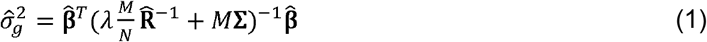

where 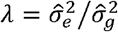 and 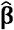 and 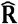 are unbiased estimates of two statistics of the population from which samples were drawn (Methods). Specifically, they are estimates of **β** the multiple regression coefficients of standardized genotypes, i.e., the additive genetic effects, and **R** the matrix of correlations between genotypes at different genetic variants, i.e., a linkage disequilibrium matrix. They are unbiased estimates under the common assumption of i.i.d. sampling from the population, and do not rely on any further assumptions about, for example, the genetic architecture of a phenotype. It is worth re-emphasising, that they are estimates of these parameters of the sampling population even if this population does not follow the assumed variance component model. This means 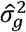 can be directly contrasted with the true additive genetic variance captured by the chosen set of genetic variants, which can be expressed in terms of **β** and **R** as

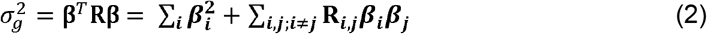

(Methods). Here, the explicit decomposition highlights the two terms contributing to 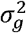 which are the genic variance, which is always positive, and the contribution due to LD between genetic variants, which can be either positive or negative.

While the presence of λ in (1) means that the provided expression does not yield an explicit solution for 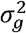, it is interpretable and provides information about the asymptotic value of 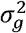 as well as it’s qualitative behavior in the finite sample setting.

Expression (1) can in general be used to qualitatively analyse the behavior of 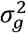 for different choices GRM and their relation to population parameters. To illustrate this point, we will in the following answer two questions. The first is, what is the expected behavior of the estimate when the standard GRM is used? The second is, what is the correct form of GRM to estimate 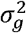?

To answer both questions we first observe by comparing (1) and (2), that 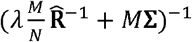 represents the effective LD structure implicitly assumed by G-REML. It depends on the choice **Σ**, and as such the GRM, and furthermore changes with the ratio of *M* and *N*. This effective LD structure does incorporate an estimate of **R**, the population’s true LD structure. However, as the sample size increases, the contribution of this term diminishes due to the presence of the *N*^−1^ factor. In particular, asymptotically as *N* increases relative to *M*, the implicit LD structure only depends, through **Σ**, on the choice of GRM.

### Estimates under the standard GRM

We can now consider the consequences of the widely used standard choice of GRM, proposed by van Raden^18^, and used as a default in popular implementations of G-REML, like GCTA^7^, BOLT-REML^9^ or DISSECT^10^. This GRM is given by 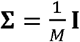. This choice will asymptotically estimate 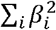, the genic variance (Methods). However, for smaller sample sizes we expect the estimate to be closer to 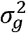 due to the stronger contribution of 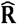. This means, that while *M*/*N* is large, if linkage disequilibrium contributes positively or negatively to 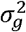, 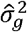 will respectively over- or under-estimate the genic variance of the chosen set of genetic variants. These conclusions are borne out by simulations (Fig. 1a, Supplementary Fig. 1, and Methods). These effects transfer to estimates of the heritability (Supplementary Fig. 2).

**Fig. 1:**
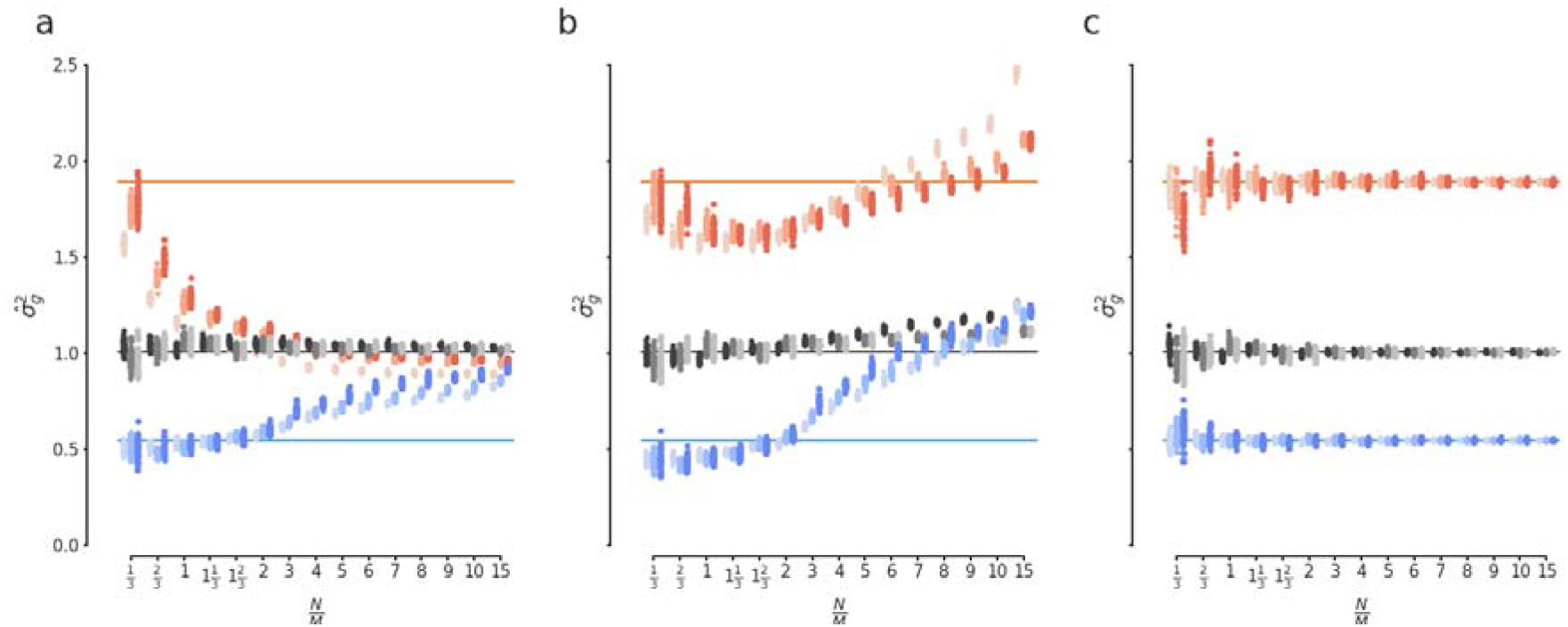
Consequences of contributions to the additive genetic variance from LD for estimates of genetic variance under different GRMs. We show G-REML estimates of the genetic variance for simulated phenotypes with either positive (red), neutral (gray) or negative (blue) contributions to the additive genetic variance from LD for varying ratios of the sample size and the number of genetic variants in the model. Different shades correspond to three different values of. Point correspond to individual estimates in replicated simulations, lines indicate true simulated values for the genetic variance. The used GRMs are **(a)** the standard GRM, **(b)** the LDAK GRM, and **(c)** the LD structured GRM. In all cases the results follow the theoretical predictions.

The described effects manifest in available real data and, as we illustrate, can lead to misleading inference. Using height data from the UK Biobank^19^ we evaluate the behavior of the estimate of 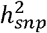 of common genetic variants for increasing sample sizes (Methods). This is consistent with a positive contribution from LD to the captured additive genetic variance (Fig. 2a). As such, we would expect estimates of heritability at a given sample size to increase when the number of genetic variants in the model increases, i.e., the ratio *N*/*M* decreases. However, addition of variants to the model could also increase SNP heritability estimates by capturing additional genetic variance through better tagging of causal variants. In order to disentangle these two effects, we increase the number of variants by adding genotypes permuted amongst individuals, which by their very nature should not capture genetic variance (Methods). As predicted, the estimates of heritability for a fixed sample size increase when the number of genetic variants modeled is doubled or quadrupled (Fig. 2b). These increases are consistent with our expectations, as becomes apparent when the results are plotted as a function of the ratio *N*/*M* (Fig. 2c). As can be seen on this scale, the added permuted genetic variants do not capture meaningful additional variance.

**Fig. 2:**
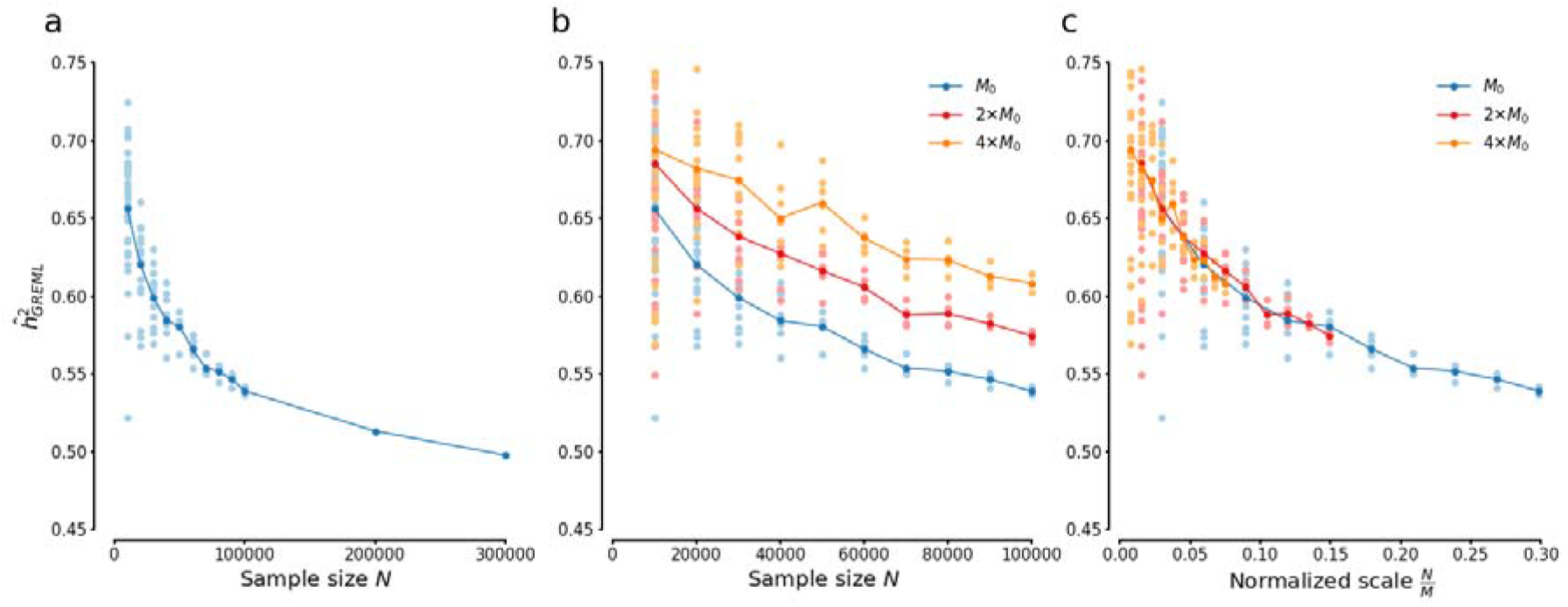
Estimates of heritability of height using a standard GRM. Estimates of heritability captured by common genetic variants for height in white-british UK Biobank individuals with varying sample sizes and numbers of genetic variants. For each sample size we obtained estimates from non overlapping subsamples of all available individuals. **(a)** Estimates heritability and their variation as a function of the sample size for a fixed set of common genetic variants. The plot shows individual estimates and their mean. **(b, c)** Heritability estimates for changing numbers of genetic variants as mutliples of, plotted as either a function of samples size or a function of the ratio of sample size and numbers of genetic variants included in the model. Additional variants beyond are generated by permuting genotypes amongst individuals. Plot show individual estimates and the means for different.

In order to further illustrate how the described effects can lead to misleading conclusions, we consider the question how much additional heritability of height is captured by rarer variants (Methods). Using 100,000 individuals the estimate of 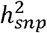 for common variants (MAF > 5%) is 0.53 (s.e. 0.005). Including rarer variants (MAF > 1%) increases this estimate to 0.66 (s.e. 0.006). However, the latter model contains almost twice the number of genetic variants significantly altering the ratio *M*/*N*. Including the genotypes for the same rarer genetic variants, but permuted amongst individuals should not increase the captured heritability. However, the estimate with permuted variants is 0.64 (s.e. 0.006). That is, a vast majority of the increase in heritability supposedly captured by rarer variants can be attributed to the change in *M*/*N*. These observations replicate across three disjoint samples of individuals and, with reduced effect, if an equivalent number of permuted common, rather than rarer, genetic variants is used (Methods and Supplementary Table 1).

### Consistent estimation of 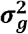

As the standard choice of GRM does not lead to a consistent estimate of 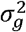, is there a GRM that does? As asymptotically 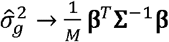 (Methods), we see by reference to (2) that the only generally correct choice of GRM is given by 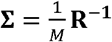, considering that *a priori* we do not know which elements of **β** have non-zero values. Although we do not know **R**, it can be consistently estimated from genotype data. An important consequence is, that the required GRM does not depend on knowledge of the underlying genetic architecture of the phenotype. This result is again borne out in simulations (Fig. 1c). This corresponds to the LD corrected GRM, recently independently proposed as an alternative to the standard GRM based on empirical observations^13^.

### Consequences of the nonrandom distribution of causal variants with respect to LD

Understanding the relationship of 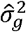 and the actual population parameters allows us to formally address questions which previously could only be considered using simulations. This is important as simulations by necessity only consider a finite set of conditions. This may lead to false conclusions if the set of considered conditions is too narrow. To illustrate this point, we turn to the question of the consequences of biases in the distribution of causal genetic variants. It has been observed that causal variants are not expected to consist of a random sample from all genetic variants^20^. Furthermore, the non-random distribution of causal variants with respect to LD in particular has been repeatedly suggested as a cause of bias in G-REML^4,8,12^. This has given rise to a number of variants on the original method, in particular LDAK^8^ and GREML-LDMS^4^, aimed at correcting this bias. Here, we re-evaluate these claims through the perspective of (1).

We will denote by 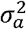 the total genetic variance of causal variants and by 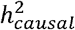 the associated SNP-heritability. We now consider estimation using a set of genetic variants which does not include the causal variants. The additive genetic variance captured by these variants, denoted by 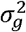, will be smaller than 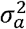, and thus 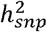 for these variants will be lower than 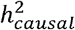.

Using the LD corrected GRM, i.e., a GRM with 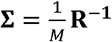, will yield estimates of 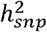. This means estimates will be smaller than 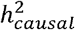. The magnitude of this difference will depend on the strength of LD between the causal variants and those in the model; increasing as LD weakens. We refer to this measure of LD as the tagging of the causal variants. In contrast to the LD corrected GRM, the behavior of estimates under other GRMs will not only depend on 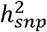, but also on the architecture of 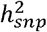. If 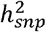 contains a positive or negative contribution of LD, estimates may change with sample size. In this context 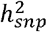 is expected to contain contributions from LD, even if the causal variants themselves are in linkage equilibrium. This is the case as multiple genetic variants which are in LD with a single causal variant will have non-zero additive effects, i.e., *β*’s, and may be expected to also be in LD with each other. We illustrate these points in the context of a previously proposed simulation study^8^. We simulate phenotypes with a constant 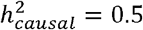 0.5, but based on casual variants which differ in how well they are tagged by the modeled genetic variants (Methods). As expected, the captured genetic variance, 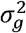, increases for phenotypes with the strength of tagging of causal variants, but always remains lower than 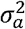, the total genetic variance of the causal variants (Fig. 3a). Crucially, the composition of 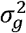 changes depending on the strength of tagging (Fig. 3a). For very weakly tagged causal variants we observe a large negative contribution from LD to 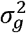. On the other hand, for very strongly tagged variants we observe a positive contribution of LD. At the same time, the genic component of 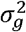 diminishes as the strength of tagging increases. The contributions of LD to 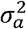 for all sets of causal variants are negligible (Supplementary Fig. 3). This means, that we do not need to resort to processes like assortative mating, that induce LD between causal variants. Even in the absence of any such process, in a typical analysis contribution of LD may be expected to play an important role.

**Fig. 3:**
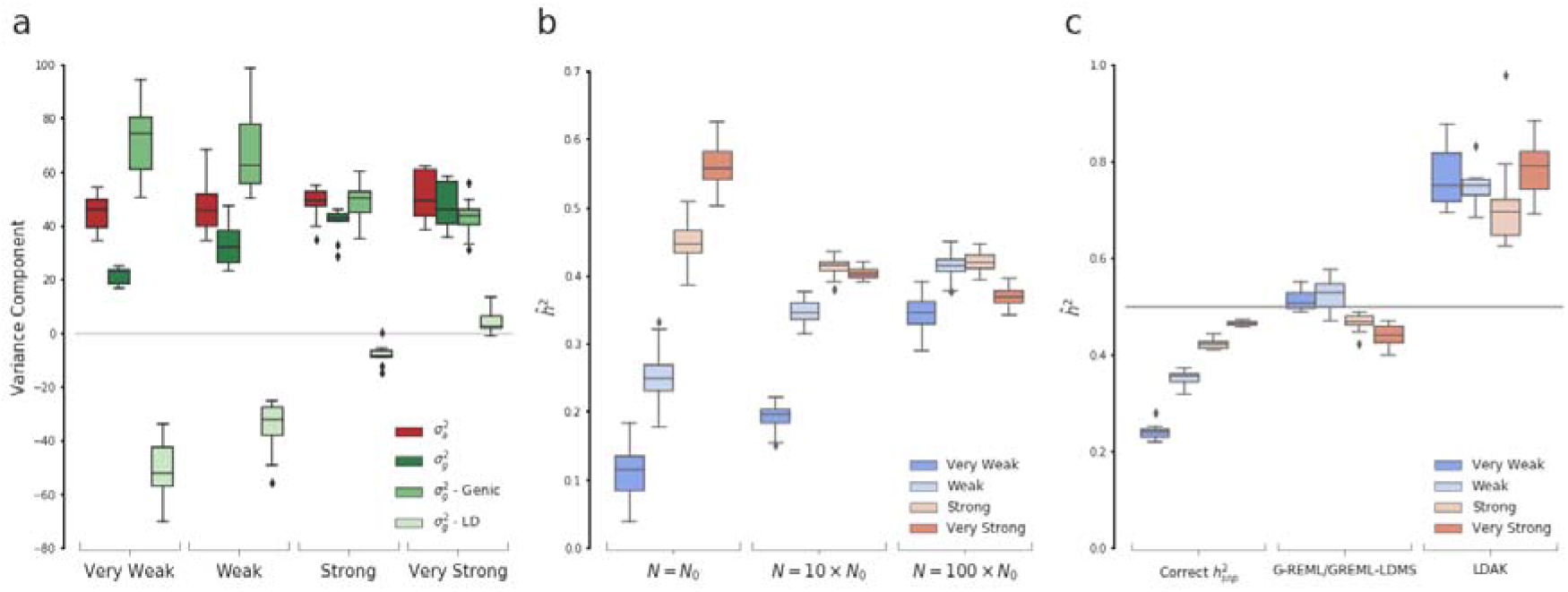
Consequences the non-random distribution of causal variants with respect to LD. **(a)** The architecture of, the additive genetic variance captured by a set genetic variants, as a function of the tagging of the underlying causal variants. **(b)** Estimates of heritability obtained using G-REML with a standard GRM for different sample sizes. **(c)** Comparison of the simulated heritability of causal variants, indicated by the gray line, the true and the expected asymptotic estimates for various G-REML methods. All plots summarise 50 replications of the simulations, with center line, box limits, whiskers and points indicating median, interquartile range, first and last datum in 1.5x interquartile range, and outliers respectively.

Use of the LD corrected GRM yields estimates which are stable across sample sizes and consistent with 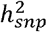 (Supplementary Fig. 4c). Estimates obtained using G-REML with the standard GRM show the expected behavior based on the observed architecture of 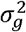 (Fig. 3b). Specifically, for small sample sizes we qualitatively replicate the original results of the simulations. That is, estimates increase with strength of tagging, and for very strongly tagged causal variants not only over-estimate 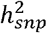, but even the full heritability 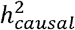. However, the results change dramatically as we increase the sample size, no longer supporting conclusions drawn based on results in smaller samples. In particular, all estimates under-estimate 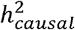 independent of the strength of tagging with biases being comparable.

In simulations, both LDAK and GREML-LDMS have been observed to reduce the bias relative to 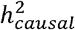 when *M*/*N* is large, i.e., comparable to the presented simulations for small sample sizes^8,14^. Using our results, we can evaluate the robustness of these empirical observations with respect to changes in *M*/*N* by considering the asymptotic behavior of these methods (Fig. 3c). The asymptotic behavior of GREML-LDMS is particularly interesting. As this approach is based on stratification of genetic variants into multiple variance components, equation (1) is not directly applicable. However, extending (1) to a multiple variance component setting, we show that asymptotically, when the standard GRM is used, the estimates of any stratified model are the same as that of the single component model containing all variants (Methods). That is to say for large sample sizes, relative to the number of genetic variants in the model, GREML-LDMS will yield the same estimates as G-REML with one standard GRM. Overall the asymptotic results for both methods, lead us to conclude that they do not represent robust approaches to estimation of either 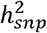 or 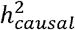. We do not question that both methods may yield good results for a specific range of *M*/*N*, but do not see a practical way to ensure a particular analysis falls within this regime. This conclusion is borne out in practice when applying LDAK for increasing sample sizes (Supplementary Fig. 4b).

## Discussion

We provide a qualitative understanding of the behavior of G-REML estimates obtained using different GRMs. We resolve any questions about the relationship of the G-REML estimates and parameters of the sampling population. Crucially, we do so without requiring any assumptions about the genetic architecture or other properties of the population. We use these insights to illustrate conditions under which G-REML estimates can be misleading. An important implication of our results is, that comparison of G-REML estimates for the same trait across analyses which differ in sample size or the numbers of genetic markers used is arguably not meaningful. Such comparisons are often made inadvertently within a single analysis. For example, modelling categories of genetic variants using multiple variance components, will involve comparisons of estimates for components potential comprising very different numbers of genetic variants^6^.

While we have centered our discussion primarily on the most widely used GRM, the standard GRM, similar analyses can be performed and confirmed in simulations for other GRMs that have been proposed in the literature. One may be inclined to argue in favor of a particular GRM based on its asymptotic behavior. We caution against this for two reasons. For one, the majority of analyses to date have been performed far from the asymptotic regime, i.e., *N*/*M* < 1. In particular, use of imputed or whole genome sequencing genotype data likely entails very low *N*/*M* ratios. Based on such data it has been recently proposed that application of G-REML-LDMS can capture the entire narrow sense heritability of human height^5^. Based on the results presented here we would predict that this conclusion will change as sample sizes increase consistent with Fig. 2a. We would further predict that, as sample sizes increase, the observed differences between G-REML-LDMS and G-REML with one variance component, which has been observed to severely overestimate the SNP heritability for dense genotype data^14^, will disappear. The second reason is that care has to be taken with the interpretation of the asymptotic estimate. It is tempting to interpret the estimate as the SNP heritability in a population with a LD structure given by **Σ**. For example one might suggest that the genic variance, as estimated asymptotically under the standard GRM, leads to the SNP heritability in a hypothetical founder population which is in linkage equilibrium. This view is in general not coherent as the estimate is based on additive effects *β* in a population which exhibits a different LD structure. To illustrate this point, we may consider estimates in the setting when causal variants are not included in the model. Here the asymptotic estimates are non-zero, because the modeled variants capture part of the effects of causal variants through LD, as can be seen in the analysis of tagging (Fig. 3c). In contrast the corresponding SNP heritability in a population which is in complete linkage equilibrium is zero, due to the lack of LD between modeled and causal variants.

As we demonstrate no complex processes acting on LD like, for example, assortative mating or selection, are necessary to give rise to complex behavior of the estimator. It is sufficient for the causal variants to not be included in the model, and most analyses may be expected to fall within this setting. Our simulations suggest that the consequences are particularly severe if causal variants are weakly tagged, which again constitutes the expected norm rather than exception. In general we may expect that the overall contribution from LD is the result of the confluence of several independent processes. Some of these processes may depend on the specific set of model variants chosen, as is for example the case for the tagging structure.

We only present an overview of some of the implications of our theoretical results. However, we think that the presented results are sufficient to warrant a critical evaluation of previous conclusions based on G-REML estimates and of claims about the efficiency of various variants of G-REML based methods. On a final note, taking a broader perspective, we anticipate that estimating the additive genetic variance directly from estimates of **β** and **R** will prove to be a more successful approach than G-REML^1,15^, by virtue of providing more flexibility to incorporate prior knowledge.

## Methods

### Derivation of Theoretical Result

#### Definition of 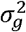 and 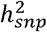

The narrow sense heritability of a phenotype in the population, *h*^2^, is defined as the ratio of 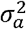, the additive genetic variance, and the phenotypic variance, 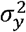 ^16,21^. In the context of individuals genotyped at a fixed set of genetic variants, the SNP heritability, 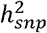, is given by the ratio of the captured additive genetic variance, 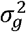, to 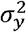 ^2^.

The additive genetic variance is defined as the variance of additive genetic values, also referred to as breeding values in the context of animal breeding, in a population of interest. It plays an important role in quantitative genetics, arising as a parameter in expressions for various quantities of interest^16^.

Formally the additive genetic value is defined by means of linear regression of genetic values on the genotype, where genetic values are given by the expected phenotype conditional on a fixed genotype^21^. Specifically, we consider a set of *M* genetic variants in the population, which, for simplicity, we shall assume are none redundant and bi-allelic. We then denote by *g* the vector of counts of one of the two alleles, chosen arbitrarily, for each of the *M* variants. The additive genetic value of a phenotype *y* with respect to the set of chosen genetic variants for an individual *i* is then defined as

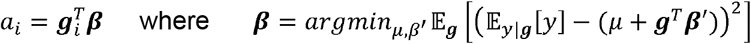

where 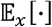 denotes the expectation w.r.t. *x*. It is worth emphasising that the formulation of *α* does not suppose a causal mechanism, in particular *β* are not the causal effects of a genetic variant, rather *α* represents expected differences between individuals from the population of interest carrying different alleles. The additive genetic variance captured by the *M* chosen variants is then defined as the variance of *α* in the population of interest^21^. Consequently it takes the form

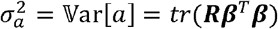

where 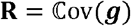 is the covariance matrix of the genetic variants in the population of interest.

#### Genomic Linear Mixed Models and G-REML

The basic Genomic Linear Mixed Model (GLMM) underlying G-REML takes the form

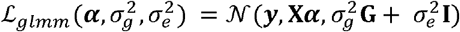

where ***y*** is a vector of phenotypes, **X** is a matrix of covariates with associated effect vector ***α***, **G** is a relationship matrix which is assumed to be positive semi-definite, and 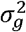, 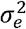 are the variance parameters of interest^3^.

Restricted maximum likelihood estimation (REML) provides an approach for unbiased inference on 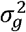, 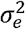. It has been classically formulated as maximum likelihood inference utilising a set of error contrasts rather than the observations themselves^21,22^. Specifically, note that **H**_**X**_ = **1** − **X(X**^*T*^**X**)^−1^**X**^*T*^ is normal and has *N* − *K* nonzero eigenvalues, where *K* is the column rank of **X**. Hence the eigen decomposition of **H**_**X**_ can be written as **L**^*T*^**L**, and **L**^*T*^**LX** = 0 and **LL**^*T*^ = **1**. The error contrast are defined by ***y***′ = **L*y*** with likelihood function

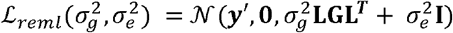

maximization of which yields the the G-REML estimates, 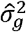, 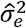.

#### General form of 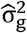

The G-REML problem, i.e., maximization of 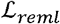, is equivalent to

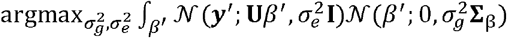

where **U** ϵ ℝ^*N*−*K*×*M*^ and **Σ**_β_ ϵ ℝ^*M*×*M*^ are a matrix with full column rank and a positive definite matrix respectively, such that **LGL**^*T*^ = **UΣU**^*T*^. We note that the latter decomposition is not unique, but does always exists as **G** was assumed to be positive semi-definite. We now observe that, up to factors independent of *β*′,

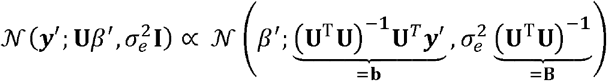

and hence,

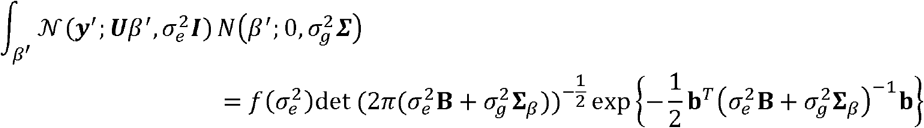

for an appropriate *f*(·). While maximization of this form with respect to 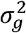 as a function of 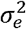 is not any more tractable than that of 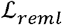, we may consider the equivalent constrained maximisation problem,

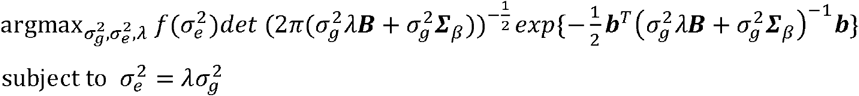

This transformation is well defined provided 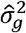, 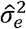 do not lie on the boundary of the parameter space, i.e., both are finite and non zero. Furthermore, 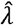 is asymptotically bounded provided both 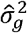 and 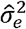 are asymptotically bounded. Considered as a function of 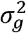, the objective is of a form proportional to the density function of a scale inverse chi squared distribution with parameters

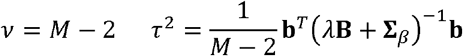

and, as a function of λ, 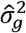 is given by the mode of said distribution. Specifically,

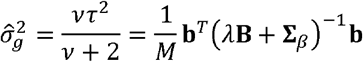

#### Form of 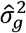 for common GRMs

We now turn to the specific case **G** = **ZΣZ**, where **Z** is a matrix of genotypes at *M* genetic variants under some encoding and is some given symmetric positive definite matrix, discussed in the Results section.

We note that, ***U*** = **LZ**, 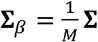 represents a suitable decomposition for the previously discussed result of the general cases to be obtained, provided **LZ** has full column rank, which may be expected to be the case once for sufficiently large sample sizes (we require at least *N* + *K* > *M*). Therefore in this setting,

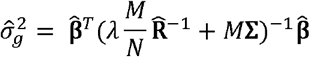

where, with **Z**_**X**_ = **H**_**X**_**Z** the genotypes with covariates regressed out, 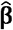is the solution of the ordinary least squares multiple regression of *y* on **Z**_**X**_ and 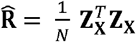, i.e., is the empirical estimate of the 2^nd^ moments of the covariate adjusted genotypes.

When furthermore, **X** = **1**, and **Z** contains standardized genotypes we recover equation in the Results section. Specifically, in this case **Z**_**X**_ = **H**_**X**_**Z** = **Z** and 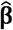,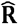 are empirical estimates of the additive genetic effects, and the matrix of correlations between genotypes at different genetic variants, i.e., a linkage disequilibrium matrix.

Considering the asymptotic setting when *N* → ∞, we observe that 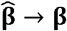, where, we recall, **β** are the population additive genetic effects. Furthermore, as 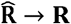. and *λ* is bounded, 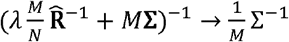, so that

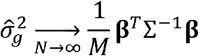

#### Asymptotic equivalence of single variance component and stratified models

The general behavior of models with multiple genomic variance components is more complicated due to dependencies between the parameters, similar to the dependency between 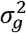 and 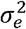 in the single genomic variance component model. However, the situation simplifies in the asymptotic regime as *N* → ∞ for constant numbers of genetic variants. In this setting we outline the argument why stratifying genetic variants into multiple variance components does asymptotically not differ from fitting a single genomic variance component, when standard GRMs are used.

We consider the case of *J* genomic variance components, where each component *j* is given by a standard GRM computed from a set of genetic variants *S*_*j*_, such that all *S*_*j*_ are disjoint. Following similar steps as for the single component model, we observe that the G-REML estimate such a model is given by

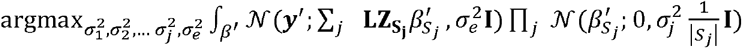

where 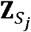 denotes the subset of genotypes at variants *S*_*j*_. We may now note that the first term under the integral is unchanged from the single genomic component setting and in particular

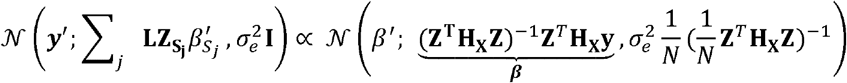

where ***β*** is the vector of population coefficient in the regression of genotypes on phenotypes adjusted for covariates. This means, in the asymptotic setting,

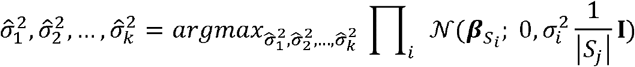

where 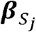 is the subvector of ***β*** at variants in *S*_*j*_. As all factors only depend on a single 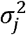, the problem decomposes into *J* independent maximizations. In particular the solution is given by 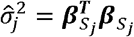. Summing over the variance components, this is the genic variance, the same quantity obtained asymptotically in a model with a single variance component including all genetic variants, i.e., the union of all *S*_*j*_.

### Data

#### UK Biobank Data

All simulations and primary data analyses were performed using data from the UK Biobank^19^, in particular the same set of genotype data was used throughout. These were genotypes of 343,884 unrelated (Kinship Coefficient < 0.0442) White-British individuals. We only considered bi-allelic autosomal variants which were assayed by both genotyping arrays employed by UK Biobank, passed UK Biobank quality control procedures and, in the unrelated White-British sample, had a minor allele frequency >1% and did not depart from Hardy-Weinberg equilibrium (P < 10^−50^). The unrelated White-British subset of individuals was obtained by excluding individuals who were identified by the UK Biobank as outliers based on either genotyping missingness rate or heterogeneity, or whose sex inferred from the genotypes did not match their self-reported sex. We then identified a subset of individuals such that for no two individuals the Kinship Coefficient was larger than 0.0442. The White-British subset of these was obtained by retaining all individuals for whom the projection on the 20 leading genomic principal components was within three standard deviations from the mean of all individuals who self identified as White-British. Finally, we removed individuals with a genotype call missing-ness rate >5% across variants which passed our quality control procedure.

#### Computation of Genomic Relationship Matrices

We make use of three types of GRMs, referred to as the standard, LDAK, and LD structured GRM. Denoting by **Z** the *N* × *M* matrix of standardized genotypes all three GRMs take the form **G** = **ZΣZ**^*T*^ for an appropriate **Σ**. Specifically, for the standard GRM 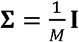. In particular, we note that as in BOLT-REML we normlise by *M*, unlike the GRM computed by GCTA which accounts for missing genotype variants in each pair of individuals. For LDAK, **Σ** is a diagonal matrix with *diag*(**Σ**) = ***w***/Σ_*m*_***w***_*m*_ where ***w*** is a vector of weights as described by Speed et al.^8^. Rather than computing weights for each sample, we computed weights once for each set of genetic variants employed in an analysis using all 343,884 available individuals. The weights were computed using the LDAK5 software. For the LD structured GRM, 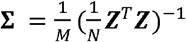, that is the empirical LD matrix. As in the case of LDAK, we computed this matrix only once for each required set of variants using all available individuals.

### Simulations

We implemented two simulation studies to illustrate aspects of the analytical results. All simulations utilized the genotype data from the UK Biobank as described above. However, we only used a restricted set of genetic variants, specifically only those on chromosome 18, in order to be able to achieve a wider range of ratios between numbers of individuals and genetic variants in the models.

All models were fitted using either GCTA, for smaller sample sizes and non-standard GRMs, or BOLT-REML, in the case of sample sizes larger than 100,000, tools.

#### Consequences of non-zero contributions from LD to genetic variance

We aimed to highlight the behavior of estimates of *h*^2^ as the ratio of *N* and *M* changes if linkage disequilibrium contributes to the additive genetic variance. To this end, we simulated phenotypes using sets of causal variants selected so that linkage disequilibrium between these variants would make a positive, neutral, or negative contribution to the additive genetic variance. For each scenario we generated 10 replicate phenotypes. We then estimated variance components using different GRMs for increasing sample sizes *N* ranging from 3,395 to 40,729 individuals. We repeated this procedure for three different numbers of variants included in the models, by dropping subsets of non-causal variants. The casual variants were always included in the model.

Results were obtained using the 10,182 common (MAF > 5%) genetic variants on chromosome 18. We simulated phenotypes by selecting a set of 100 causal variants and giving each an effect size of 1 on the scale of normalized genotypes. In order to obtain positive, neutral, or negative contributions to the additive genetic variance from linkage disequilibrium we sampled the causal variants in pairs as follows. We repeatedly selected a new causal variant. at random from all remaining non-causal variants. Conditional on *i*, we then selected a second new causal variant *j* at random from all remaining non causal variants for which *r*_*ij*_ ∈ (*r*_*min*_,*r*_*max*_, where *r*_*ij*_ is the correlation between variants *i* and *j* computed using all individuals. We set (*r*_*min*_,*r*_*max*_) to (0.7,1.0), (−10^−5^, 10^−5^) and (−0.5,−0.4) in order to generate a positive, neutral, and negative contribution from LD respectively. The environmental effect for each individual was sampled from a normal distribution with variance chosen so that the asymptotic estimate of *h*^2^ using the standard GRM would be either 0.25, 0.5, or 0.75.

It is worth emphasizing, that we specifically chose to retain the causal variants in the model and did not simulate effects sizes following a reasonable prior distribution. We retained the causal variants so as to exclude effects of tagging of these by the variants retained in the model. We did not simulate the effects sizes from any of the linear mixed model prior distributions to illustrate the point that the presented results hold without assumptions on the true generating distribution for the population.

#### Consequences of biased distribution of causual variants

We implemented a simulation study following the design of Speed et al.^8^. Specifically, we simulated phenotypes selecting causal variants based on how well they were tagged by other variants. The measure of tagging for a genetic variant *i* was computed as 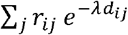, where 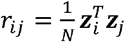 is the empirical correlation between variants *i* and *j* computed using all available individuals, *d*_*ij*_ is the distance in base pairs between these variants and, following Speed et al., *λ* = −log(0.125)/30^6^. We denoted variants with tagging values from the bottom 0-20% and 20-40% of values amongst all considered variants as *very weakly* and *weakly* tagged. Similarly, variants with a value from the top 0-20% and 20-40% are denoted as *strongly* and *very strongly* tagged. It is worth noting that this definition differs from Speed et al. who denoted the whole bottom and top 40% as weakly and strongly tagged respectively. We prefer to consider non-overlapping ranges, but as a consequence our results differ slightly from those of Speed et al. even in the small sample setting. Specifically, we do not observe overestimation of the true heritability for the strongly tagged causal variants.

Results were obtained using all 16,887 available genetic variants on chromosome 18. We used 50 causal variants. We performed 50 replications. We fitted models using the standard GRM for one hundreds (N=N_0_=3,438), one tenth (N=34,388) and all (N=343,884) available individuals.

### Application to Height in UK Biobank

We estimated heritability using the standard G-REML approach for height measured in UK Biobank participants. We included Sex, Age and the leading five genomic principle components, computed on the whole UK Biobank cohort, as covariates in the analysis.

#### Effect of changes in sample size

We fitted models for different sample sizes, using all 334,942 available common (MAF>5%) genetic variants. For each sample size *N* we randomly divided the available 343,884 individuals into the maximal number of non-overlapping subsets, i.e., [343,884 / *N*] subsets, and fitted models for each subset. The number of available estimates therefore decreases with *N*, down to only a single estimate for *N* = 200,000 and *N* = 300,000.

#### Effect of changes in number of variants

In order to study the effect of varying the number of variants without changes to the SNP heritability we generated genotype data which in addition to the *M*_0_ = 334,942 available common (MAF>5%) genetic variants included variants which by design do not contribute SNP heritability. To this end, we combined *k* copies of the original genotype data for the common genetic variants, to obtain genotypes with *k* · *M*_0_ genetic variants. In each but the first copy of the genotypes the genotypes of all variants were jointly randomly permuted amongst individuals. This ensures, that the resulting genotypes maintained the statistical characteristics of the original genotypes, amongst others, the MAF frequency spectrum, distribution of LD. Crucially, the true underlying SNP heritability is not affected by the addition of permuted genotypes as these are uncorrelated with the phenotype.

In addition to results for the original genotype data (*k* = 1), we fitted models for different sample sizes, using generated genotype data for *k* = 2,4. We again divided the available 343,884 individuals into the maximal number of non-overlapping subsets for each sample size and fitted models for each subset.

#### Variance captured by rarer variants

We fitted models using either all available genetic variants with either MAF > 5%, *M* = 334,942, or MAF > 1%, *M* = 623,496, for three disjoint subsets of 100,000 individuals each. We then obtained genotypes with *M* = 623,496 variants but with the same expected captured additive genetic variance in two ways. On the one hand, we permuted the genotypes for genetic variants with MAF <5% and >1% between the individuals in each subset. We permuted all genetic variants jointly, i.e., an individual *i* was given the genotype for these variants from one individual *j*. We combined these permuted genotypes with the un-permuted genotypes for variants with MAF > 5%. In the second approach, we sampled 288,554 of the variants with MAF > 5%, permuted the genotypes for these amongst individuals and added them as new variants to the un-permuted genotypes of variants with MAF>5%. With then fitted the models using these two sets of partly permuted genotypes.

## Accession codes

This research has been conducted using the UK Biobank Resource under project 788.

## Code availability

Models were fitted using GCTA v1.24.7 and bolt-LMM v2.3. LDAK v5.0 was used to compute weights for the LDAK GRM.

## Acknowledgements

This research has been conducted using the UK Biobank Resource under project 788. The work was funded by the Roslin Institute Strategic Programme Grant from the BBSRC (BB/P013732/1). A.T. also acknowledges funding from the Medical Research Council and O.C.-X. from MRC fellowship MR/R025851/1.

