## Supplementary Figures and Tables for "SNP heritability: What are we estimating?"

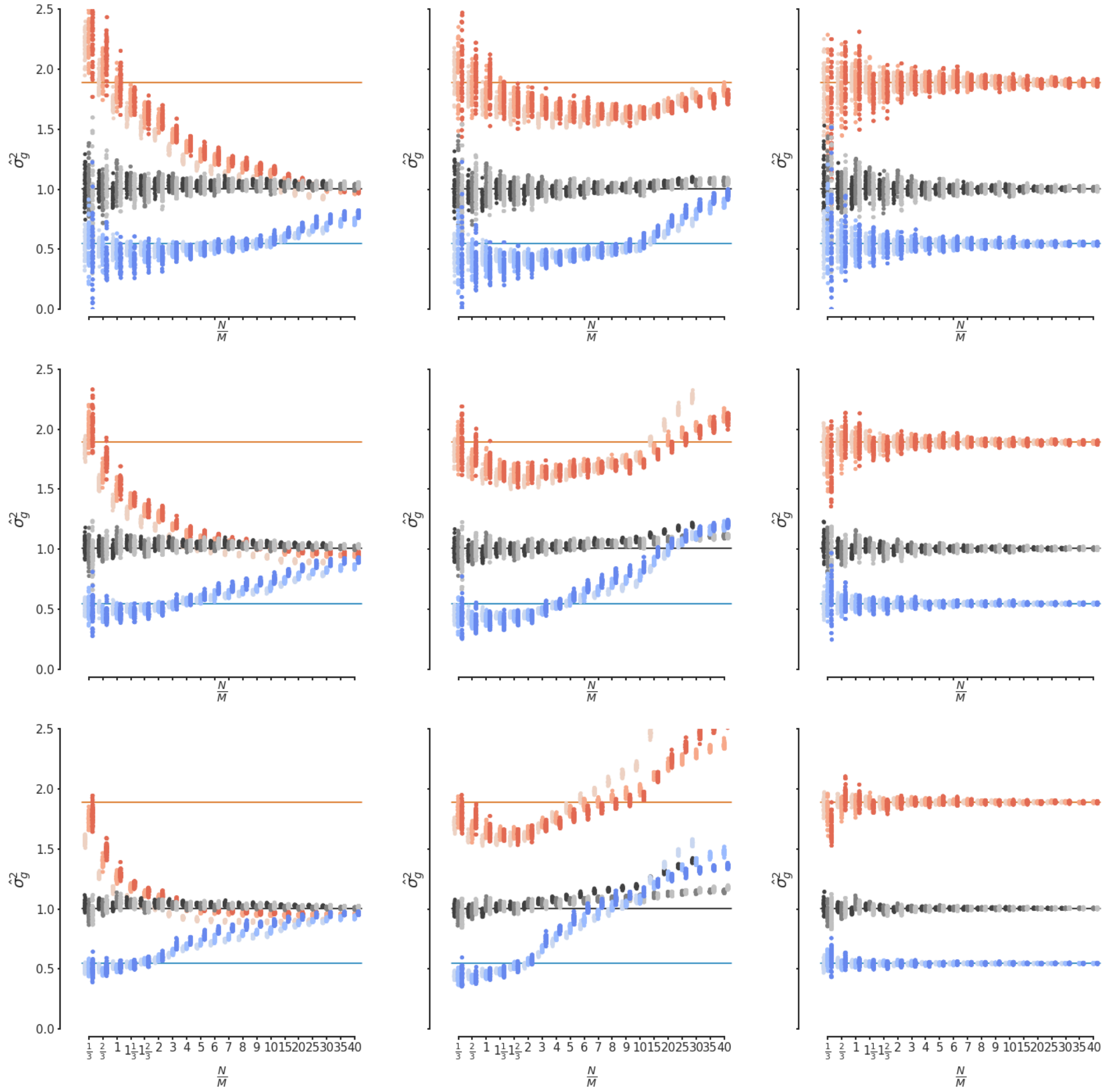


**Supplementary Fig. 1: Consequences of contributions to the additive genetic variance from LD for estimates of genetic variance under different GRMs and heritabilities.** We show G-REML estimates of the genetic variance for simulated phenotypes with either positive (red), neutral (gray) or negative (blue) contributions to the additive genetic variance from LD for varying ratios of the sample size $N$ and the number of genetic variants in the model $M$. Different shades correspond to three different values of $M$. Point correspond to individual estimates in replicated simulations, lines indicate true simulated values for the genetic variance. Each column corresponds to a different GRM with **(left column)** the standard GRM, **(middle column)** the LDAK GRM, and **(right column)** the LD structured GRM. Each row corresponds to a different value of residual variance, chosen to yield heritabilities of **(top row)** 0.25, **(middle row)** 0.5, and **(bottom row)** 0.75 in the case of zero contribution from LD. The middle row corresponds to Fig. 1 in the main text.

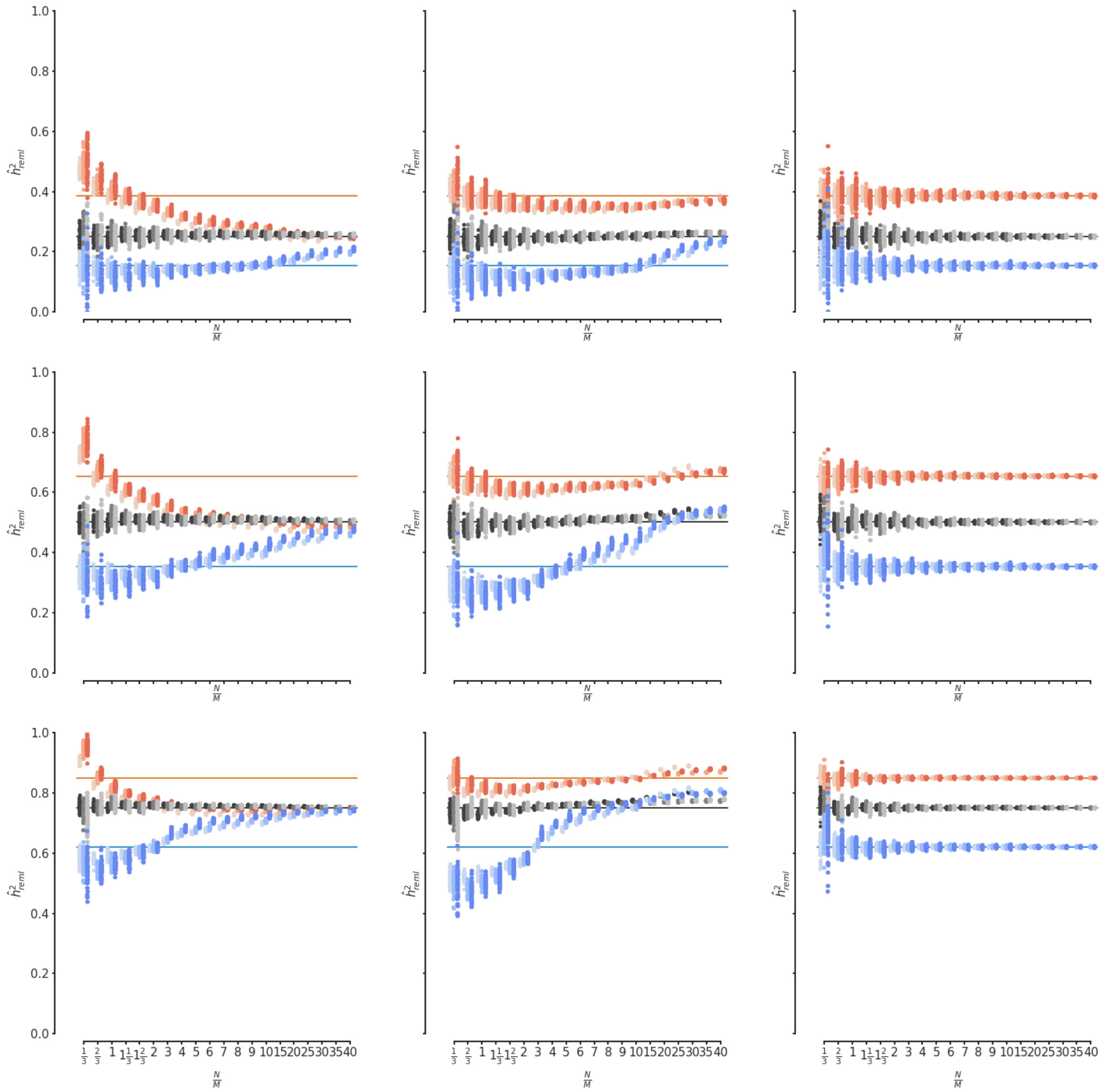


**Supplementary Fig. 2: Consequences of contributions to the additive genetic variance from LD for estimates of heritability under different GRMs and heritabilities.** Results of the same analyses as in Supplementary Fig1, but showing heritabilities. That is, we show G-REML estimates of the heritability for simulated phenotypes with either positive (red), neutral (gray) or negative (blue) contributions to the additive genetic variance from LD for varying ratios of the sample size $N$ and the number of genetic variants in the model $M$. Different shades correspond to three different values of $M$. Points correspond to individual estimates in replicated simulations, lines indicate true simulated values for the heritability. Each columns corresponds to a different GRM with, **(left column)** the standard GRM, **(middle column)** the LDAK GRM, and **(right column)** the LD structured GRM. Each row corresponds to a different value of residual variance, chosen to yield heritabilities of **(top row)** 0.25, **(middle row)** 0.5, and **(bottom row)** 0.75 in the case of zero contribution from LD.

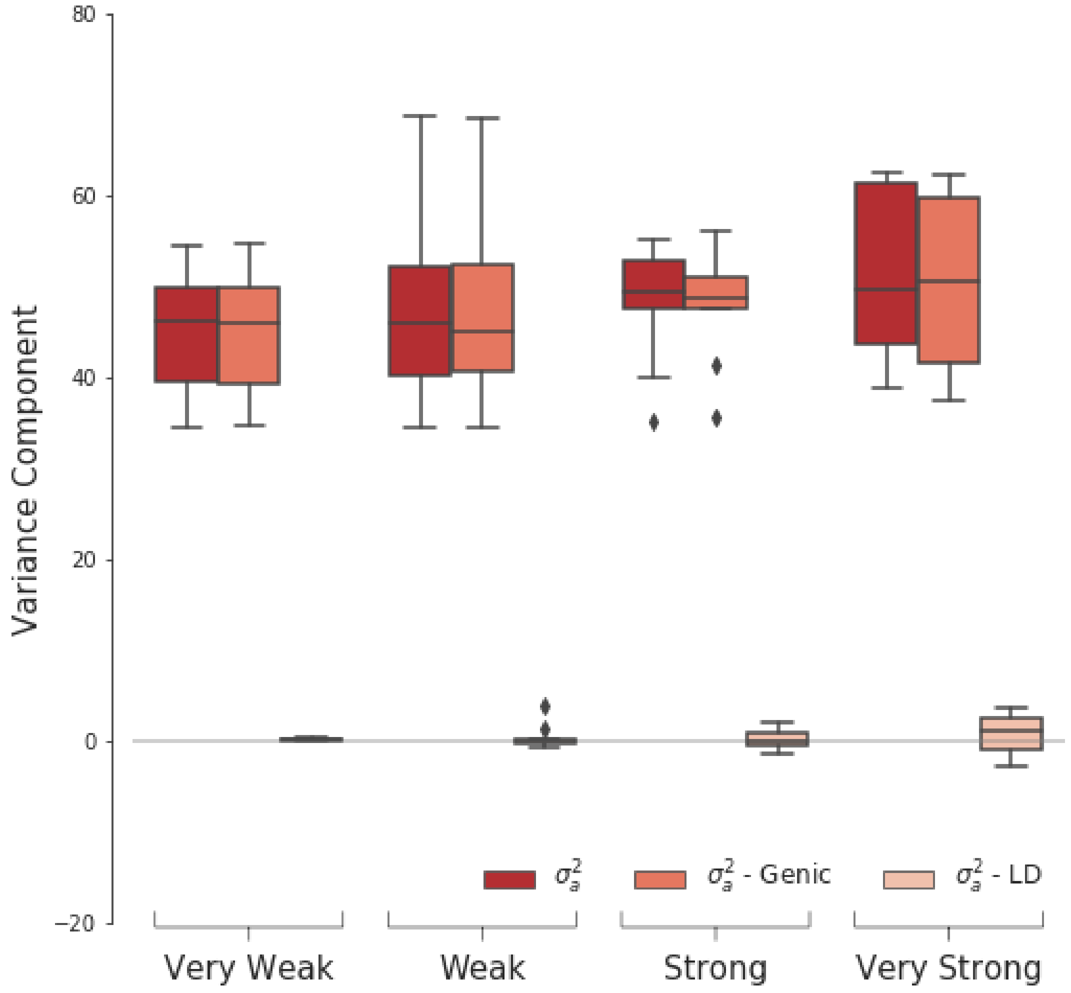


**Supplementary Fig. 3: Decomposition of simulated genetic variance of Fig. 3 in the main text, into genic and LD contribution component.**

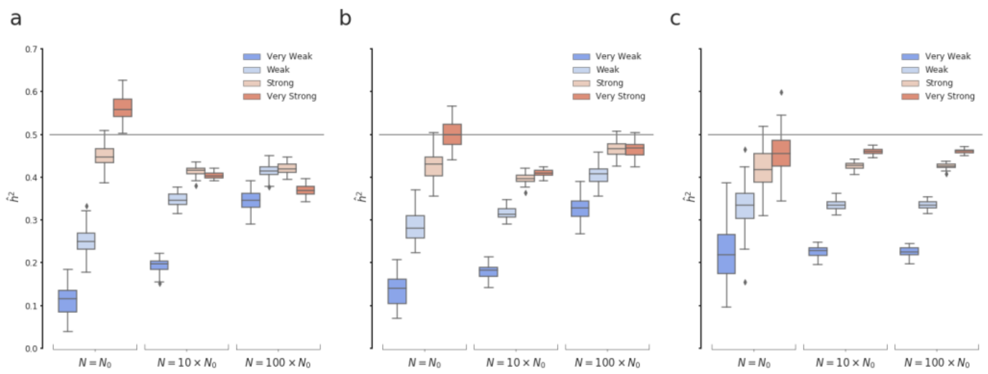


**Supplementary Fig. 4: Consequences the non-random distribution of causal variants with respect to LD on heritability estimates under different GRMs.** Estimates of heritability obtained using G-REML with a standard GRM for different sample sizes for simulated phenotypes with different tagging of the underlying causal variants. Plots correspond to different choices of GRM, with **(a)** the standard GRM, **(b)** the LDAK GRM, and **(c)** the LD structured GRM. Plot **(a)** is equivalent to Fig. 3b in the main text and reproduced here for completeness.

**Supplemetary Table 1: Heritability of height captured by combinations of common and rare genetic variants for 100,000 UK Biobank participants.**

| Variants in model† | $\hat{h}^2_{reml}$ (s.e.) | | |
|---|---|---|---|
| | Replicate 1 | Replicate 2 | Replicate 3 |
| Common | 0.53 (0.005) | 0.53 (0.005) | 0.54 (0.005) |
| Common + Rare | 0.66 (0.006) | 0.66 (0.006) | 0.68 (0.006) |
| Common + Permuted Common | 0.57 (0.006) | 0.57 (0.006) | 0.58 (0.006) |
| Common + Permuted Rare | 0.64 (0.006) | 0.64 (0.006) | 0.65 (0.006) |

†Fitted models contained one standard GRM computed from a combination of non-permuted and permuted common (MAF ≥ 5%) and rare (5% > MAF > 1%) genetic variants.
